# Exploring the Genetics of Lesion and Nodal Resistance in Pea (*Pisum sativum* L.) to *Sclerotinia sclerotiorum* Using Genome-wide Association Studies and RNA-Seq

**DOI:** 10.1101/233288

**Authors:** Hao-Xun Chang, Hyunkyu Sang, Jie Wang, Kevin E. McPhee, Xiaofeng Zhuang, Lyndon D. Porter, Martin I. Chilvers

## Abstract

The disease white mold caused by the fungus *Sclerotinia sclerotiorum* is a significant threat to pea production and improved resistance to this disease is needed. Nodal resistance in plants is a phenomenon where a fungal infection is prevented from passing through a node and the infection is limited to an internode region. Nodal resistance has been observed in some pathosystems such as the pea (*Pisum sativum* L.)-*S. sclerotiorum* pathosystem. Other than nodal resistance, different pea lines display different levels of stem lesion size restriction, referred to as lesion resistance. It is unclear whether the genetics of lesion resistance and nodal resistance are identical or different. This study applied genome-wide association studies (GWAS) and RNA-Seq to understand the genetic makeup of these two types of resistance. The time series RNA-Seq experiment consisted of two pea lines (the susceptible ‘Lifter’ and the partially resistant PI 240515), two treatments (mock samples and *S. sclerotiorum* inoculated samples), and three time points (12, 24, and 48 hours post-inoculation). Integrated results from GWAS and RNA-Seq analyses identified different redox-related transcripts for lesion and nodal resistances. A transcript encoding a glutathione S-transferase was the only shared resistance source for both phenotypes. There were more leucine rich-repeat containing transcripts found for lesion resistance, while different candidate resistance transcripts such as a VQ motif-containing protein and a myo-inositol oxygenase were found for nodal resistance. This study demonstrated the robustness of combining GWAS and RNA-Seq for identifying white mold resistance in pea, and results suggest different genetics underlying lesion and nodal resistance.

## 1 INTRODUCTION

*Sclerotinia sclerotiorum* (Lib.) de Bary the causal agent of white mold disease, is one of the most destructive plant pathogens worldwide. *S. sclerotiorum* is capable of infecting more than 400 host plants and causes millions of dollars of crop yield losses each year (Bolton, Thomma, & Nelson, 2006). Several studies have reported different secondary metabolites, effectors, and pathogenicity factors of *S. sclerotiorum* that are involved in establishing the infection (Bolton, Thomma, & Nelson, 2006; Mbengue et al., 2016; Wei & Clough, 2016). One of the well-known virulence strategies is the production of oxalic acid, which creates a low pH and acidic environment for infection (Xu et al., 2015). Oxalic acid suppresses reactive oxygen species (ROS) produced by plants at the beginning of infection and generates a reducing status that favors colonization (Williams et al., 2011). Fine-tuned redox homeostasis from the initial reducing status to the later oxidative status in plant tissues is important for *S. sclerotiorum* to switch from the initial hemibiotrophic lifestyle to the later necrotrophic lifestyle (Kabbage, Yarden, & Dickman, 2015). Studies searching for plant resistance to *S. sclerotiorum* have found quantitative interactions (McCaghey et al., 2017), and potential resistance genes included those with functions to maintain ROS and redox stresses during *S. sclerotiorum* infection (Girard et al., 2017; Seifbarghi et al., 2017; Zhou, Sun, & Xing, 2013).

Pea (*Pisum sativum* L.) is an important legume crop in the United States, and white mold continuously causes substantial damage and yield reduction (Tayeh et al., 2015). Similar to soybean, *S. sclerotiorum* infection primarily begins when ascospores of *S. sclerotiorum* colonize blooms and invade through petioles into the stem. Severely infected plants will wilt and lodge. Resistance to white mold in pea has been observed via two different phenotypes, the first is lesion size where the length of stem lesion is measured after inoculation. The second phenotype is referred to as nodal resistance, and appears to be a unique mode of resistance which has been observed in some varieties of pea and soybean (Calla et al., 2009; Porter, Hoheisel, & Coffman, 2009; Porter, 2011). Nodal resistance can be defined as the inhibition of lesion expansion at a node limiting pathogen colonization of plant stem tissue. Restriction of lesion expansion at the nodes has also been observed for stem-infecting fungi such as *Diaporthe* and *Macrophomina* species on soybean and cowpea (Hobbs, Schmitthenner, & Ellett, 1981; Muchero et al., 2011). However, nodal resistance has been rarely documented and other than knowing lignin content is negatively correlated with nodal resistance in soybean (Peltier et al., 2009), our understanding is limited.

Transcriptomics and differential expression (DE) analysis using RNA-Seq have become a standard approach to identifying resistance genes for white mold, and many studies have applied this approach to oilseed rape (*Brassica napus*) (Girard et al., 2017; Seifbarghi et al., 2017; Zhuang et al., 2012). While most of these studies focused on the expression comparisons between a resistant and a susceptible variety, the genetic diversity of white mold resistance in *B. napus* might be underestimated using only this approach. Genome-wide association study (GWAS) is a robust approach to map white mold resistance and to capture the resistance diversity in a germplasm collection (Moellers et al., 2017; Wei et al. 2016; Wei et al. 2017). The GWAS approach has been demonstrated in soybean (*Glycine max* Merr. L.) resistance to *S. sclerotiorum* where numerous SNPs associated with this quantitative resistance were discovered (Bastien, Sonah, & Belzile, 2014; Moellers et al., 2017; Wu et al., 2016a). However, mapping results may discover single nucleotide polymorphisms (SNPs) that locate in intergenic genomic regions, and the interpretation of a confidence interval relies on the size of linkage disequilibrium (Bush & Moore, 2012). RNA-Seq and GWAS both have their advantages, but combined they provide a powerful tool to discover not only active genes that express in response to treatments, but also genetic diversity and SNPs associated with the treatment. This combined strategy has been applied to understand white mold resistance and yields in *B. napus* (Lu et al., 2017; Wei et al., 2016) but not in pea. Because genes that can be found by both GWAS and RNA-Seq will have higher potential in contributing to white mold resistance, this study aimed to understand and compare the genetics of lesion and nodal resistance by applying both GWAS and RNA-Seq approaches in the pea-*Sclerotinia sclerotiorum* pathosystem.

## 2 MATERIALS AND METHODS

### 2.1 GWAS: data source and analysis

Data used for GWAS was published in Porter, Hoheisel, & Coffman (2009). Briefly, there were 282 pea lines with a mean lesion resistance rating. The white mold fungus (*Sclerotinia sclerotiorum*) Scl02-05 isolated from pea in Quincy, Washington, USA in 2003 was used for inoculations (Porter, Hoheisel, & Coffman, 2009). A mean lesion resistance rating was obtained after 72 hpi in a humid greenhouse and day/night temperature ranges around 28°C /15°C. There were 266 pea accessions with nodal resistance ratings. Nodal resistance was measured using an ordinal scale from 0 to 5 after two weeks post inoculation, where 0 = dead plant; 1 = lesion expanded down the stem from the fourth inoculated node to the first node; 2 = lesion expanded from the fourth to the second node; 3 = lesion expanded from the fourth node to the third node; 4 = lesion did not expand beyond the initial inoculation point at the fourth node (Porter, Hoheisel, & Coffman, 2009). There were four to eight replications to represent each accession. The USDA Pea Single Plant Plus Collection with single nucleotide polymorphism (SNP) data included in this study (Holdsworth et al., 2017). Association test was conducted in PLINK version 1.9 (Purcell et al., 2007). Population stratification was controlled using a pairwise identity-by-state (IBS) clustering with a maximum clustering node of 2 and a *p* value cutoff of 0.05 for the pairwise population concordance test. The IBS clustering matrix was included in a basic association test, and a minor allele frequency (MAF) of 0.05 was applied. The empirical *q* value at 0.01 from an adaptive permutation test with default parameters was used to determine association significance. The GBS raw reads containing significant SNPs were searched against the Trinity *de novo* transcriptome (assembled in the following sections) using BLASTN to acquire annotations at an E value cutoff of 10^-5^.

### 2.2 Plant inoculations for RNA-Seq

A white mold-susceptible pea (*Pisum sativum* L.) cultivar ‘Lifter’ (PI 628276) and a white mold-partially resistant pea accession, PI 240515, were used in this study. The same *Sclerotinia sclerotiorum* isolate (Scl02-05) used to generate the GWAS data was used for RNA-Seq experiments. Seeds from ‘Lifter’ and PI 240515 were planted at a depth of 1 cm in pasteurized soil in a plastic pot (approximately 170 cm^3^). The soil consisted of a mixture of 85 L of Special Blend Soil Mix (Sun Gro Horticulture, Bellevue, WA), 113 L of propagation-grade coarse perlite (Supreme Perlite Company, Portland, OR), and 900 g of Scotts Osmocote Classic 14-14-14 (The Scotts Company, Marysville, Ohio). Plants were grown in a growth chamber at 23°C/20°C (day/night), with a photoperiod of 14 h and 170 μmol quanta (s^-1^m^-2^) for two weeks. One day before *Sclerotinia sclerotiorum* inoculations, pea plants were covered with a thick transparent plastic cover, which filtered the amount of light reaching the plants down to 45-55 μmol quanta (s^-1^m^-2^), and maintained a high humidity (RH %; 86.91 ± 13.45; WatchDog 1000 Series, Spectrum Technologies Inc., Aurora, IL). Pea plants were inoculated at the fourth node leaf axil with a 49 mm^3^ *S. sclerotiorum* colonized agar plug from the leading edge of a culture grown on potato dextrose agar (PDA; BD Company, Sparks, MD). Mock inoculations were performed with sterile PDA plugs.

### 2.3 RNA extraction and sequencing

The RNA-Seq experiment had a time series factorial design with two varieties (‘Lifter’ and PI 240515), two treatments (mock and *S. sclerotiorum* inoculation), and three time points at 12, 24 and 48 hours post inoculation (hpi). For each condition, two biological replicates of pea samples were collected. In order to acquire RNA samples that provide both expression data for lesion and nodal resistance, tissues within 2 cm of the inoculated fourth node were collected from at least 12 plants for each biological replicate. These tissues were immediately frozen in liquid nitrogen. Total RNA was isolated using Trizol^®^ reagent (Invitrogen, CA) according to manufacturer’s instructions. DNase digestion (Promega, WI) was performed on the RNA extract to remove potential DNA contamination. RNA samples were further purified using the RNeasy Plant Mini Kit (Qiagen, Valencia, CA) and quality verified using a 2100 Bioanalyzer RNA Nanochip (Agilent, Santa Clara, CA). Samples achieved a RNA integrity number (RIN) value above 7.5 and were quantified using the Qubit ^®^ 2.0 Fluorometer (Invitrogen, Carlsbad, CA) and a total of 10 μg RNA was used for cDNA library preparations following the Illumina TruSeq RNA Preparation Kit manufacturer’s instructions (Illumina, San Diego, CA). A paired-end 2×75 base sequencing was run on the Illumina GA IIx sequencer (Illumina, San Diego, CA) at the Research and Technology Support Facility at Michigan State University.

### 2.4 *De novo* transcriptome assembly

Illumina raw reads were quality checked using FastQC version 0.11.5 (Andrews, 2010) and quality controlled using FASTX-toolkit version 0.0.14 (Gordon, 2014). Reads with 90 percent length above Phred score 30 were kept for analyses. Trimmomatic version 0.33 was used to remove adapters and to separate paired reads and single reads (Bolger, Lohse & Usadel, 2014), and only paired reads were used for *de novo* assembly. All samples were pooled and aligned to the complete nearly gapless *S. sclerotiorum* genome sequence (Derbyshire et al., 2017) using the sensitive mode of Bowtie2 version 2.2.6 and Tophat2 version 2.1.0 (Kim et al., 2013). Reads unmapped to *S. sclerotiorum* genome were *de novo* assembled by Trinity version 2.4.0 using K-mer size 25, 29, and 32 (Grabherr et al., 2011; Haas et al., 2013).

### 2.5 Differential expression (DE), heatmap clustering, and gene ontology (GO) analyses

A k-mer index of 31 bp was built for the Trinity *de novo* transcriptome and paired-end reads were pseudo-aligned to the index using Kallisto version 0.43.0 with 1000 bootstrap (Bray et al., 2017). DE analysis was conducted using Sleuth version 3 in default mode using transcripts per million (TPM) normalization (Bray et al., 2017). The default filter setting was applied such that transcripts with more than 5 estimated counts in 47 percent of samples were kept for DE analysis. The principal component analysis (PCA) was used to visualize variation structure among all samples, and the hierarchical clustering using Ward’s criterion. D2 method was applied to group transcripts in the heatmap analysis (Murtagh & Legendre, 2014). A time series model with three explanatory variables, including the variety (‘Lifter’ and PI 240515), the treatment (mock and *S. sclerotiorum* inoculation), and the time (12, 24, and 48 hpi), were included in a full model whereas a reduced model excluded a variable of interest. A model comparison using likelihood ratio test was used to identify transcripts with DE in response to the variable of interest, and a multiple comparison-corrected *q* value at 0.05 was used to determine the significance. *De novo* transcripts were functionally annotated by soybean coding sequences using BLASTN at an E value cutoff of 10^-5^, and soybean gene models with orthologous *de novo* transcripts of pea were subjected to agriGO v2.0 singular enrichment analysis (SEA) using Fisher’s exact test with Yekutieli correction to control false discovery rate (FDR) at 0.05 in multiple-tests (Tian et al. 2017).

## 3 RESULTS

### 3.1 Phenotype data for lesion and nodal resistance

There were 282 and 266 pea germplasm lines screened for lesion and nodal resistance, respectively. While the lesion resistance distribution approximated a normal distribution (Fig. 1a), the nodal resistance distribution was highly skewed toward zero with only 12 germplasm lines rated with a score above 3 (Fig. 1b). The lesion resistance and nodal resistance have a significant negative correlation (Pearson’s correlation coefficient: -0.19, *p* < 0.05) due to the inverse scales, meaning higher value of lesion resistance and lower value of nodal resistance represented the susceptibility. The mild correlation indicated the possibility of different genetic mechanisms of these two types of resistances (Fig. 1c). The white mold susceptible line ‘Lifter’ and the white mold partially resistant line PI 240515 were selected for RNA-Seq. PI 240515 displays not only a slower disease progress compared to the susceptible ‘Lifter’ under growth chamber conditions (Fig. 1d), but also a better resistance performance in field trials (McPhee & Muehlbauer. 2002; Zhuang et al., 2013). A time course RNA-Seq experiment was set up in a factorial design, and a total of 12 samples with about 700 million reads were acquired from Illumina sequencing (Table S1).

**FIGURE 1.**
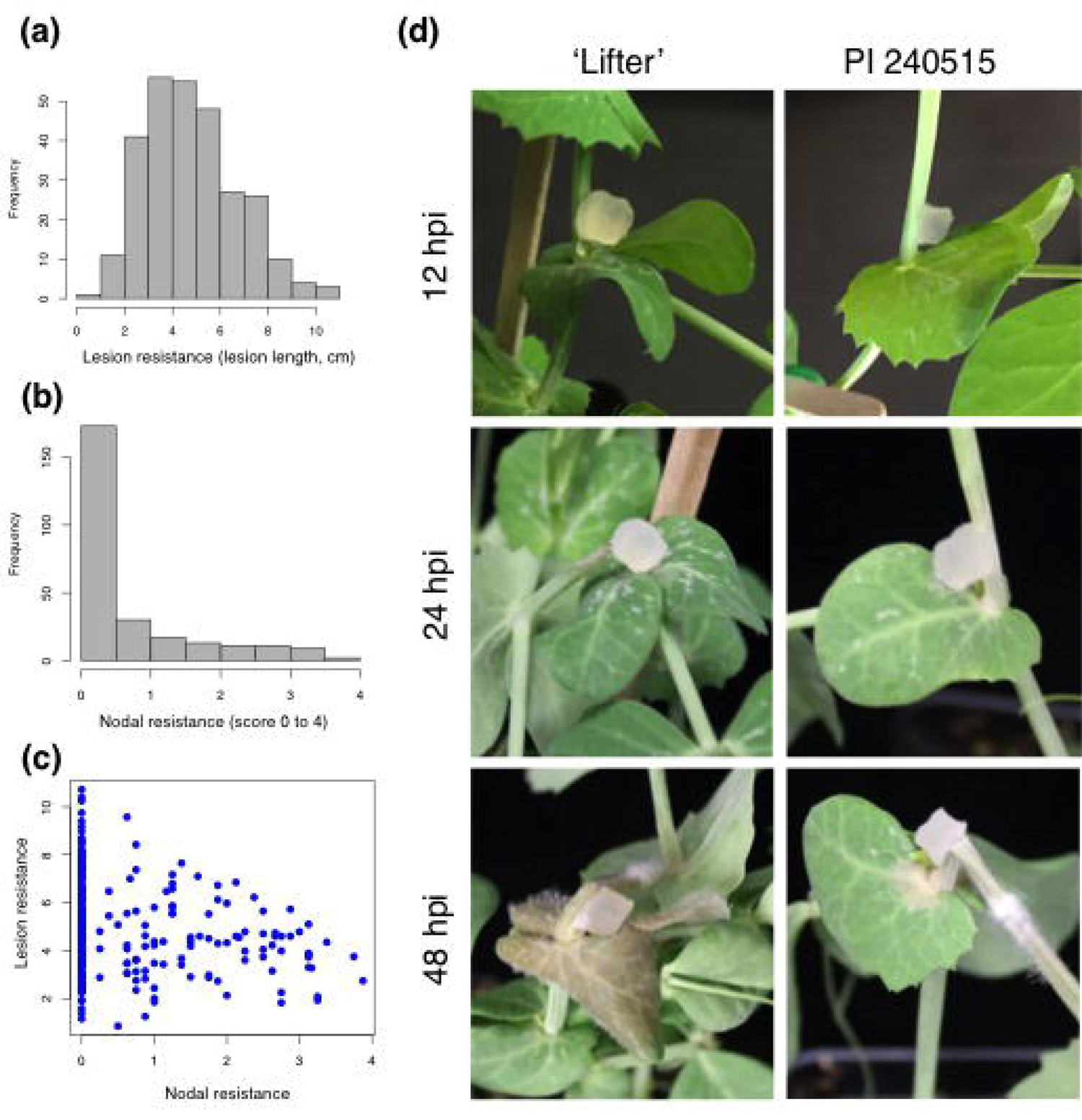
Lesion and nodal resistance phenotypes of pea lines used in GWAS, and phenotypes of ‘Lifter’ and PI 240515 used in RNA-Seq. (a) Phenotypic distribution of lesion resistance (lesion size in centimeter). (b) Phenotypic distribution of nodal resistance (score 0 = dead plant, 4 = lesion restricted to the inoculated node number 4). (c) Pearson’s correlation between lesion and nodal resistance demonstrates slight but significant negative correlation (-0.19, *p* < 0.05; negative due to the inverse rating scale for nodal resistance). (d) Phenotypic difference between a susceptible cultivar ‘Lifter’ and a partially resistant accession PI 240515 over time. A potato dextrose agar block containing actively growing hyphal tips of *S. sclerotiorum* was used for inoculation. PI 240515 has partial resistance and displays slower disease progress compared to susceptible ‘Lifter’. Infection and damping-off can be observed in ‘Lifter’ as early as 12 hpi and 24 hpi, respectively, but not PI 240515. Infection expands in ‘Lifter’ as early at 48 hpi, and infection can be observed around the inoculated site of PI 240515 at 48 hpi.

### 3.2 GWAS

A total of 35,658 SNPs were included in the association analysis using PLINK (Purcell et al., 2007). There were 206 and 118 SNPs found to be significantly associated with lesion and nodal resistance, respectively (Table S2 and S3, respectively). Without a standard genome for pea, the position and chromosome information for SNPs were deficient and it also made the annotation to these SNPs difficult. In order to understand the annotations of these significant SNPs, the original genotyping-by-sequencing (GBS) raw reads harboring each SNP were retrieved (Holdsworth et al., 2017) and searched against our RNA-Seq *de novo* transcriptome using BLASTN.

### 3.3 *De novo* transcriptome assembly

Using a stringent quality control threshold that keeps only raw reads with 90 percent of bases above Phred score of 30 (error rate 0.001%, one error per thousand bases), paired-end reads were mapped to the complete nearly gapless *S. sclerotiorum* genome using Tophat2 (Derbyshire et al., 2017; Kim et al., 2013). The *de novo* transcriptome using k-mer of 29 bp, which contains 96,588 transcripts including isoforms, resulted in the highest assembly quality (Table 1) similar to a previous *de novo* transcriptome of pea (Kerr et al., 2017), and the re-mapped rate at 80% was satisfactory based on an empirical threshold of Trinity (Grabherr et al., 2011; Haas et al., 2013). Accordingly, the k-mer 29 *de novo* transcriptome was selected and a total of 60,598 transcripts were extracted from the longest representative isoform per gene model in the k-mer 29 *de novo* transcriptome (Table 1).

**TABLE 1.**
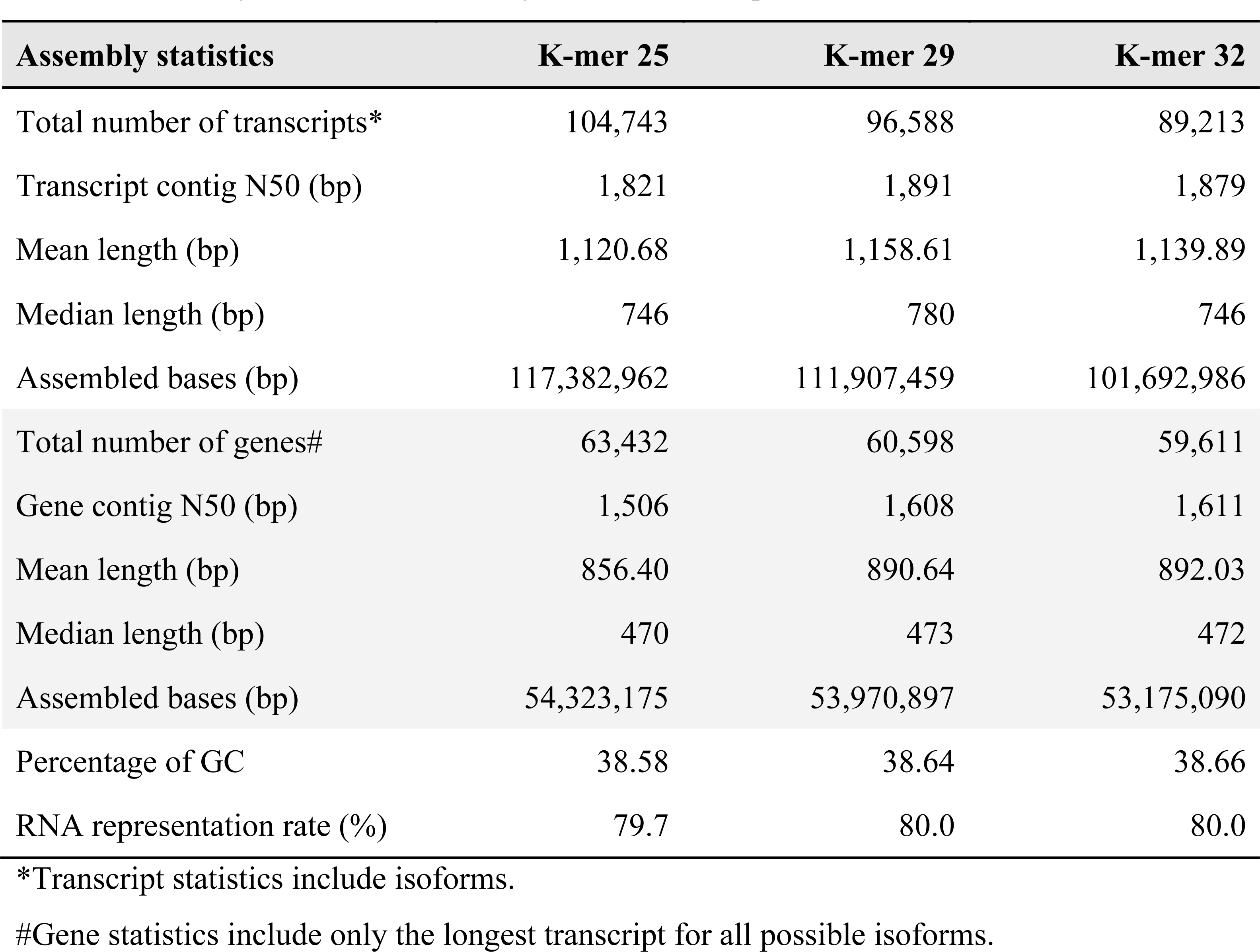
Quality assessment of Trinity *de novo* transcriptome assemblies

### 3.4 Annotation of candidate resistance genes using GWAS and RNA-Seq

There were 206 significant SNPs associated with lesion resistance, but only 96 SNPs were matched to *de novo* transcripts. Among these 96 *de novo* transcripts, 66 of them could be annotated with an orthologous soybean gene (Table 2, Table S2). In terms of nodal resistance, there were 118 significant associated SNPs, and 61 SNPs could be matched to *de novo* transcripts. Among these 61 *de novo* transcripts, 33 of them could be annotated with an orthologous soybean gene (Table 3, Table S3). In comparing the GWAS for lesion and nodal resistance, only one SNP (TP13557) can be found in both cases, and the *de novo* transcript containing this SNP was annotated as a putative glutathione S-transferase (GST) (similar to soybean gene Glyma.06G117800). The results supported the negative phenotypic correlation and suggested the genetic backgrounds of lesion and nodal resistance were different. Other potential transcripts found in GWAS with resistance functions included redox enzymes, cytochromes, leucine rich-repeat (LRR)-containing proteins, ABC transporters, armadillo (ARM) repeat-containing protein, pentatricopeptide repeat (PPR)-containing proteins, and tetratricopeptide repeat (TPR)-containing proteins (Collier & Moffett, 2009; Sharma & Pandey, 2015).

**TABLE 2.** Candidate resistance transcripts identified in GWAS for lesion resistance

| SNP | GWAS q value | Allele | MAF | *De novo* transcript | E value* | Gm.W82.a2.v1 | Annotation | Annotation E value |
| --- | --- | --- | --- | --- | --- | --- | --- | --- |
| TP104551 | 1.41E-04 | A/G | 0.108 | TRINITY_DN19399_c0_g1_i2 | 2.00E-06 | Glyma.05G228200 | Xylogalacturonan β-1,3-xylosyltransferase | 0.00E+00 |
| TP82039 | 1.22E-03 | C/T | 0.489 | TRINITY_DN22733_c0_g1_i1 | 7.00E-25 | Glyma.16G023200 | TPR-containing protein | 0.00E+00 |
| TP132492 | 1.25E-03 | C/A | 0.100 | TRINITY_DN11274_c0_g2_i1 | 2.00E-26 | Glyma.20G193300 | CC-NBS-LRR disease resistance protein | 0.00E+00 |
| TP176436 | 1.33E-03 | A/G | 0.356 | TRINITY_DN15345_c0_g1_i2 | 6.00E-19 | Glyma.19G233900 | Oxidoreductase | 0.00E+00 |
| TP192026 | 2.68E-03 | G/A | 0.185 | TRINITY_DN12885_c0_g1_i1 | 5.00E-26 | Glyma.04G159700 | UDP-arabinopyranose mutase | 0.00E+00 |
| TP58726 | 2.81E-03 | C/T | 0.145 | TRINITY_DN21727_c0_g1_i1 | 7.00E-25 | Glyma.17G090900 | ARM repeat superfamily protein | 0.00E+00 |
| TP46107 | 3.03E-03 | A/C | 0.400 | TRINITY_DN23674_c1_g2_i1 | 2.00E-26 | Glyma.03G168000 | Pleiotropic drug resistance ABC transporter | 0.00E+00 |
| 59136_75 | 3.61E-03 | A/G | 0.085 | TRINITY_DN24902_c0_g1_i1 | 5.00E-15 | Glyma.16G130700 | Serine carboxypeptidase | 0.00E+00 |
| 64833_37 | 3.94E-03 | T/C | 0.426 | TRINITY_DN21727_c0_g1_i1 | 1.00E-30 | Glyma.17G090900 | ARM repeat superfamily protein | 0.00E+00 |
| TP40425 | 3.96E-03 | C/T | 0.069 | TRINITY_DN11862_c0_g1_i1 | 7.00E-25 | Glyma.09G242500 | PPR-containing protein | 0.00E+00 |
| TP5714 | 3.97E-03 | A/G | 0.160 | TRINITY_DN8622_c0_g1_i2 | 7.00E-25 | Glyma.20G013200 | ARM repeat superfamily protein | 0.00E+00 |
| TP122891 | 4.37E-03 | C/T | 0.065 | TRINITY_DN20767_c0_g1_i1 | 7.00E-25 | Glyma.06G065000 | Cellulose synthase | 0.00E+00 |
| TP107762 | 4.65E-03 | C/G | 0.348 | TRINITY_DN21987_c1_g2_i1 | 2.00E-19 | Glyma.08G284100 | LRR-RLK | 0.00E+00 |
| TP105293 | 7.28E-03 | C/T | 0.490 | TRINITY_DN22449_c0_g1_i1 | 2.00E-26 | Glyma.13G359600 | PPR-containing protein | 0.00E+00 |
| TP14472 | 7.60E-03 | A/G | 0.108 | TRINITY_DN23601_c0_g1_i1 | 2.00E-19 | Glyma.08G160900 | ABC transporter | 0.00E+00 |
| 54792_76 | 7.91E-03 | C/G | 0.321 | TRINITY_DN23920_c0_g1_i14 | 2.00E-15 | Glyma.13G370300 | Pleiotropic drug resistance ABC transporter | 0.00E+00 |
| TP56848 | 9.05E-03 | A/C | 0.193 | TRINITY_DN23559_c0_g2_i3 | 2.00E-26 | Glyma.06G178400 | Copper amine oxidase | 0.00E+00 |
| 67025_44 | 9.21E-03 | T/C | 0.352 | TRINITY_DN29578_c0_g1_i1 | 6.00E-27 | Glyma.06G131900 | Cytochrome b5 | 1.00E-85 |
| TP184273 | 9.52E-03 | A/T | 0.055 | TRINITY_DN22193_c0_g1_i4 | 7.00E-25 | Glyma.19G004700 | ARM repeat superfamily protein | 0.00E+00 |
| TP13557 | 9.82E-03 | C/T | 0.194 | TRINITY_DN7903_c0_g1_i2 | 7.00E-25 | Glyma.06G117800 | **Glutathione S-transferase** | 0.00E+00 |

\* E value of BLASTN using a GBS read containing the significant SNP against the *de novo* transcriptome.

**TABLE 3.** Candidate resistance transcripts identified in GWAS for nodal resistance

| SNP | GWAS q value | Allele | MAF | *De novo* transcript | E value* | Gm.W82.a2.v1 | Annotation | Annotation E value |
| --- | --- | --- | --- | --- | --- | --- | --- | --- |
| 56725_68 | 3.74E-03 | G/C | 0.197 | TRINITY_DN5298_c0_g1_i1 | 3.00E-30 | Glyma.05G107600 | ACT domain repeat protein | 0.00E+00 |
| TP41550 | 4.50E-03 | A/G | 0.167 | TRINITY_DN25769_c0_g1_i1 | 7.00E-25 | Glyma.09G051900 | VQ motif-containing protein | 7.00E-39 |
| TP13557 | 4.59E-03 | C/T | 0.194 | TRINITY_DN7903_c0_g1_i2 | 7.00E-25 | Glyma.06G117800 | **Glutathione S-transferase** | 0.00E+00 |
| TP52272 | 4.64E-03 | T/C | 0.106 | TRINITY_DN23515_c1_g1_i4 | 1.00E-15 | Glyma.12G053900 | β-glucosidase | 0.00E+00 |
| TP164197 | 4.83E-03 | T/A | 0.093 | TRINITY_DN41476_c0_g1_i1 | 2.00E-26 | Glyma.08G254000 | PPR repeat-containing protein | 0.00E+00 |
| 161268_51 | 5.04E-03 | T/C | 0.095 | TRINITY_DN21874_c0_g2_i1 | 4.00E-35 | Glyma.04G029500 | RING/U-box superfamily protein | 0.00E+00 |
| 53592_14 | 6.47E-03 | C/T | 0.147 | TRINITY_DN21524_c0_g1_i1 | 4.00E-35 | Glyma.05G224500 | Myo-inositol oxygenase | 0.00E+00 |
| TP108888 | 6.96E-03 | C/T | 0.248 | TRINITY_DN21142_c0_g2_i2 | 7.00E-25 | Glyma.06G183500 | Protein kinase superfamily protein | 4.00E-63 |
| TP119499 | 7.49E-03 | T/C | 0.053 | TRINITY_DN43663_c0_g1_i1 | 2.00E-12 | Glyma.18G202100 | Calcium-dependent protein kinase | 0.00E+00 |
| TP163256 | 8.04E-03 | A/G | 0.357 | TRINITY_DN7950_c0_g1_i1 | 7.00E-25 | Glyma.07G018400 | Peroxisome-related protein | 0.00E+00 |
| 14350_13 | 8.25E-03 | A/G | 0.124 | TRINITY_DN16214_c1_g2_i1 | 2.00E-32 | Glyma.09G011200 | Cytochrome b-561 | 0.00E+00 |
| TP164952 | 9.52E-03 | T/C | 0.097 | TRINITY_DN23515_c1_g1_i4 | 2.00E-06 | Glyma.12G053900 | β-glucosidase | 0.00E+00 |
| 123212_21 | 9.66E-03 | G/A | 0.155 | TRINITY_DN21917_c0_g1_i2 | 2.00E-27 | Glyma.11G202700 | TPR repeat-containing protein | 0.00E+00 |
| 33803_4 | 9.96E-03 | C/T | 0.058 | TRINITY_DN23419_c1_g1_i6 | 4.00E-35 | Glyma.07G151800 | β-glucosidase | 8.00E-59 |

\* E value of BLASTN using a GBS read containing the significant SNP against the *de novo* transcriptome.

### 3.5 PCA of RNA-Seq samples

In order to understand the gene expression difference of the susceptible ‘Lifter’ and the partially resistant PI 240515 especially for those candidate resistance transcripts found in GWAS, the paired-end reads of each sample were pseudo-aligned to the *de novo* transcriptome containing 60,598 transcripts using Kallisto. A total of 17,220 genes with at least 5 estimated counts in 47% of samples and these transcripts were kept for PCA. The expressions were normalized using TPM approach and quantified by Sleuth (Bray et al., 2017). *S. sclerotiorum* inoculation appeared to be the strongest influential factor to explain the variations of gene expression and the first principal component explained about 75% of variance (x axis). The treatments separated samples to two different spaces. Mock samples were clustered in one spot regardless of the pea lines and the time points, meaning relatively similar expression patterns (Fig. 2). For *S. sclerotiorum*-inoculated samples, the time points appeared to be the second most influential factor as samples from the same time point grouped close to each other.

**FIGURE 2.**
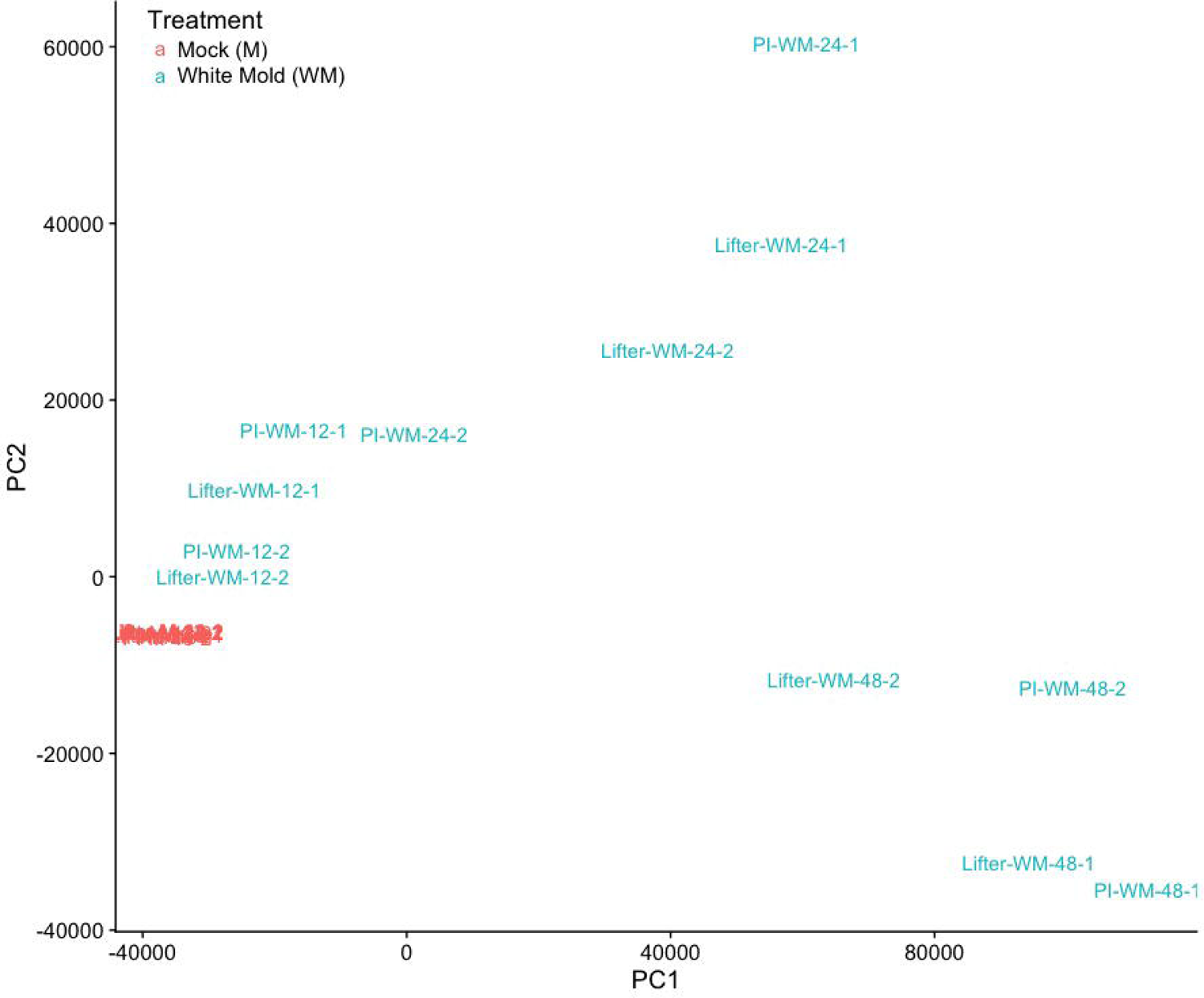
Principal component analysis. The distribution of samples are mostly influenced by the *Sclerotinia sclerotiorum* inoculation, as demonstrated by the grouping of mock inoculated samples. Principal component 1 explains about 75% of total variance from the expression based on the normalized transcripts per million.

### 3.6 DE, heatmap clustering, and GO analysis

A total of 17,220 genes were analyzed for DE using a time series model. Transcripts were clustered into four groups based on their expression patterns in the heatmap (Fig. 3). While cluster III contains 12,668 transcripts that are generally down-regulated, cluster IV contains 2,902 transcripts that are generally up-regulated in the *S. sclerotiorum* inoculated samples regardless of the pea lines. On the other hand, cluster I and II, which contain 1,506 and 954 transcripts, respectively, do not have clear expression pattern differences in *S. sclerotiorum* inoculated samples compared to mock samples. While GO analysis using the SEA approach identified general biological process, cellular component, and molecular function for transcripts in the cluster I, II, and III (Figure S1, S2 & S3), transcripts in the cluster IV were significantly enriched for oxidation reduction in the biological process (GO0055114, FDR: 7.99 × 10^-9^) and oxidoreductase activity in the molecular function (GO0016491, FDR: 3.36 x 10^-11^). These results indicated that many transcripts highly induced in cluster IV after *S. sclerotiorum* inoculation were related to redox maintenance (Fig. 4a). Other than transcripts with potential redox balancing functions, significant enrichment for transcripts with cofactor-, vitamin-, heme-, or iron-binding functions were also found (Fig. 4b). Because it has been suggested that oxalic acid stimulates iron release and soybeans were shown to express higher ferritin for capturing iron and maintaining iron homeostasis during infection (Calla et al. 2014), the enrichment results of these element-binding transcripts in pea may indicate the homeostasis struggle during infection. Although most transcripts in cluster IV had higher expression after *S. sclerotiorum* inoculation, only a few transcripts displayed significantly higher expression in PI240515 than ‘Lifter’, and together with the results from GO analysis, the possibilities that transcripts in the cluster IV are genes involved in common responses to pathogen infection could not be excluded.

**FIGURE 3.**
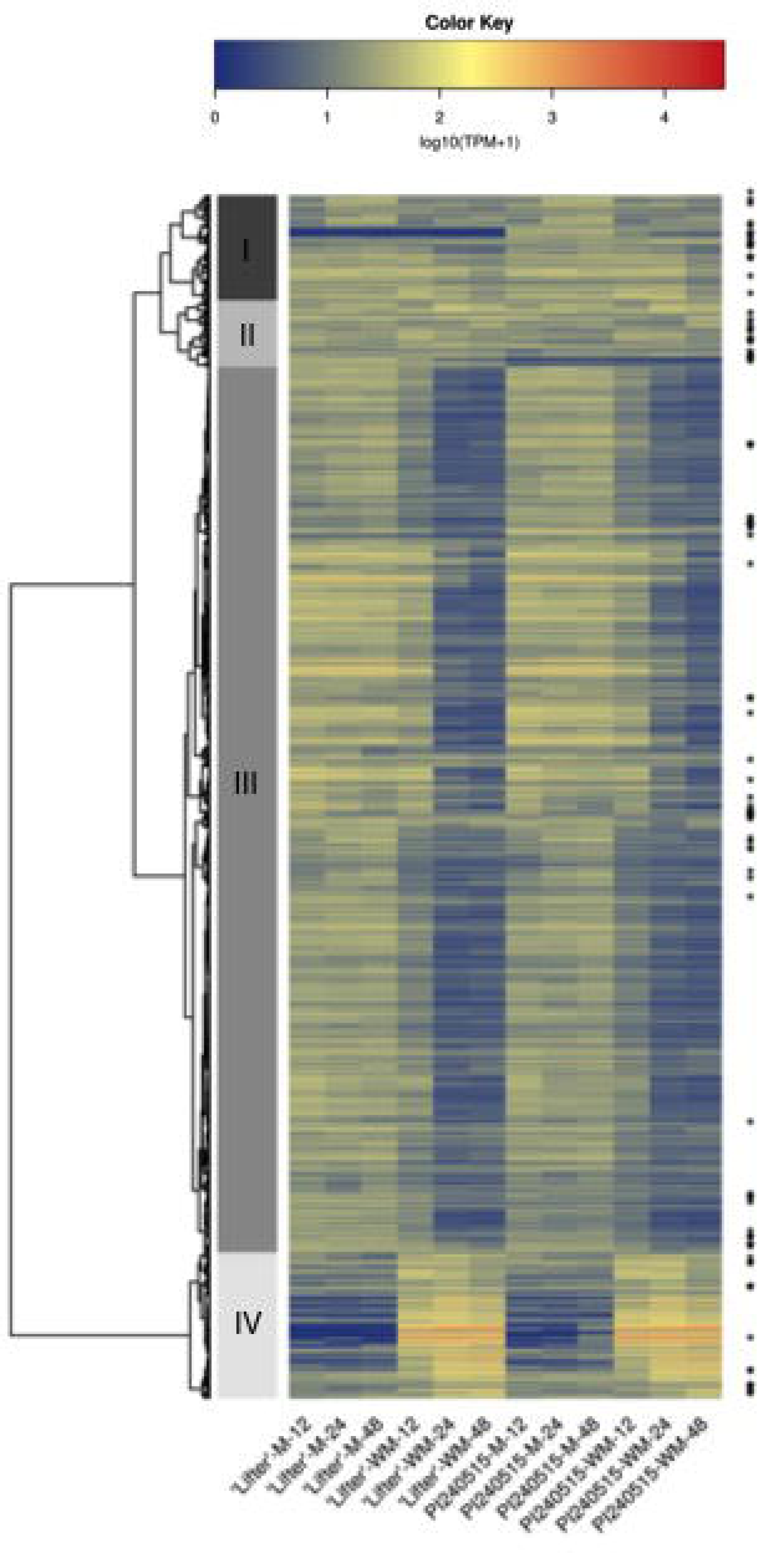
Heatmap and clustering analysis for DE transcripts over time. Transcripts with an asterisk have significant DE between ‘Lifter’ and PI 240515 only in *S. sclerotiorum*-inoculated samples but not mock samples. Clustering analysis breaks the 17,220 transcripts into four clusters. Cluster III contains transcripts that are generally down-regulated in *S. sclerotiorum* inoculated samples, and cluster IV contains transcripts that are up-regulated in *S. sclerotiorum* inoculated samples. Although cluster IV has higher expression after *S. sclerotiorum* inoculation, a few transcripts displayed significantly higher expression in PI 240515 than ‘Lifter’, indicating most of the transcripts in cluster IV may be involved in common responses to pathogen infection but not necessarily candidate resistance genes.

**FIGURE 4.**
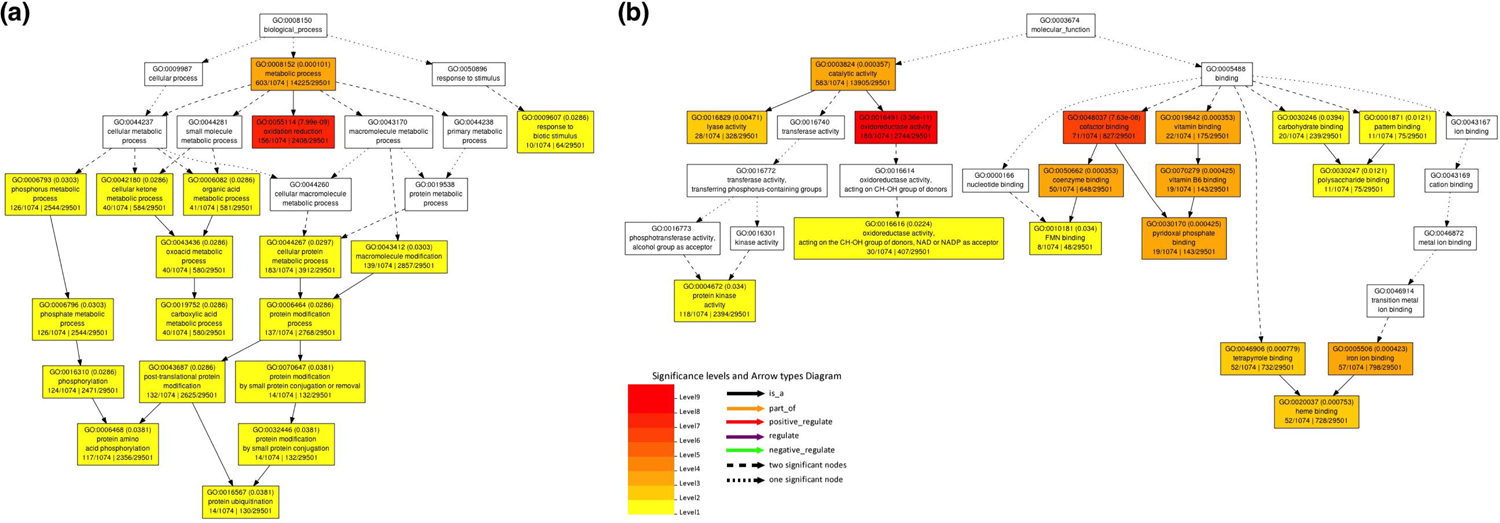
Gene ontology (GO) using singular enrichment analysis for transcripts in the cluster IV. (a) Significant GO terms in the biological process. (b) Significant GO terms in the molecular functions. Color panel shows significant enrichment from level 1 in yellow color to level 9 in red color for both (a) and (b).

In order to identify candidate genes governing *S. sclerotiorum* resistance, two assumptions were made: (i) a candidate gene should respond to *S. sclerotiorum* inoculation, and (ii) the expression of a candidate gene should have up-regulated and significant DE in the *S. sclerotiorum*-inoculated samples of PI 240515 compared to the *S. sclerotiorum*-inoculated samples of ‘Lifter’, but not the mock samples of PI 240515 compared to the mock samples of ‘Lifter’. A Venn diagram was illustrated to indicate the number of DE transcripts for four different comparisons (Fig. 5). Two of these sections fulfill our assumptions. The first contains 119 transcripts, which is the overlapping area among the purple (transcripts of ‘Lifter’ with DE in *S. sclerotiorum* inoculated samples compared to mock samples), red (transcripts of PI 240515 with DE in *S. sclerotiorum* inoculated samples compared to mock samples), and yellow blocks (transcripts of *S. sclerotiorum* inoculated samples with DE in PI 240515 compared to ‘Lifter’) but not the green block (transcripts of mock samples with DE in PI 240515 compared to ‘Lifter’) (Table S4). The second section which fulfills the assumptions contains 29 transcripts, which corresponds to the overlapping area of the red and yellow blocks (transcripts with DE in PI 240515 but not ‘Lifter’, and these transcripts had DE in PI 240515 compared to ‘Lifter’ under *S. sclerotiorum* inoculation) (Table S5). While most of these transcripts (119 + 29 transcripts) had lower expression in PI 240515 compared to ‘Lifter’ after *S. sclerotiorum* inoculation, a few transcripts had higher expression in PI 240515 than ‘Lifter’ after *S. sclerotiorum* inoculation, including three transcripts that encode LRR receptor-like kinase (LRR-RLK). The first LRR-RLK is TRINITY_DN22904_c0_g1_i2, which had nearly zero expression in mock samples, but the expressions were induced higher in PI 240515 than ‘Lifter’ after *S. sclerotiorum* inoculation (Fig. 6a). The expressions of TRINITY_DN23231_c0_g2_i2 and TRINITY_DN18054_c0_g1_i1 were higher in PI 240515 than ‘Lifter’ after *S. sclerotiorum* inoculation, but their expressions were down-regulated after *S. sclerotiorum* inoculation compared to mock samples (Fig. 6b,c). On the other hand, the expression of TRINITY_DN4777_c0_g1_i1 was generally higher in ‘Lifter’ than PI 240515, and *S. sclerotiorum* inoculation caused up-regulation more in ‘Lifter’ than PI 240525 (Fig. 6d) and the expression of TRINITY_DN21848_c0_g1_i1 was higher in mock samples than in *S. sclerotiorum* inoculated samples (Fig. 6e).

**FIGURE 5.**
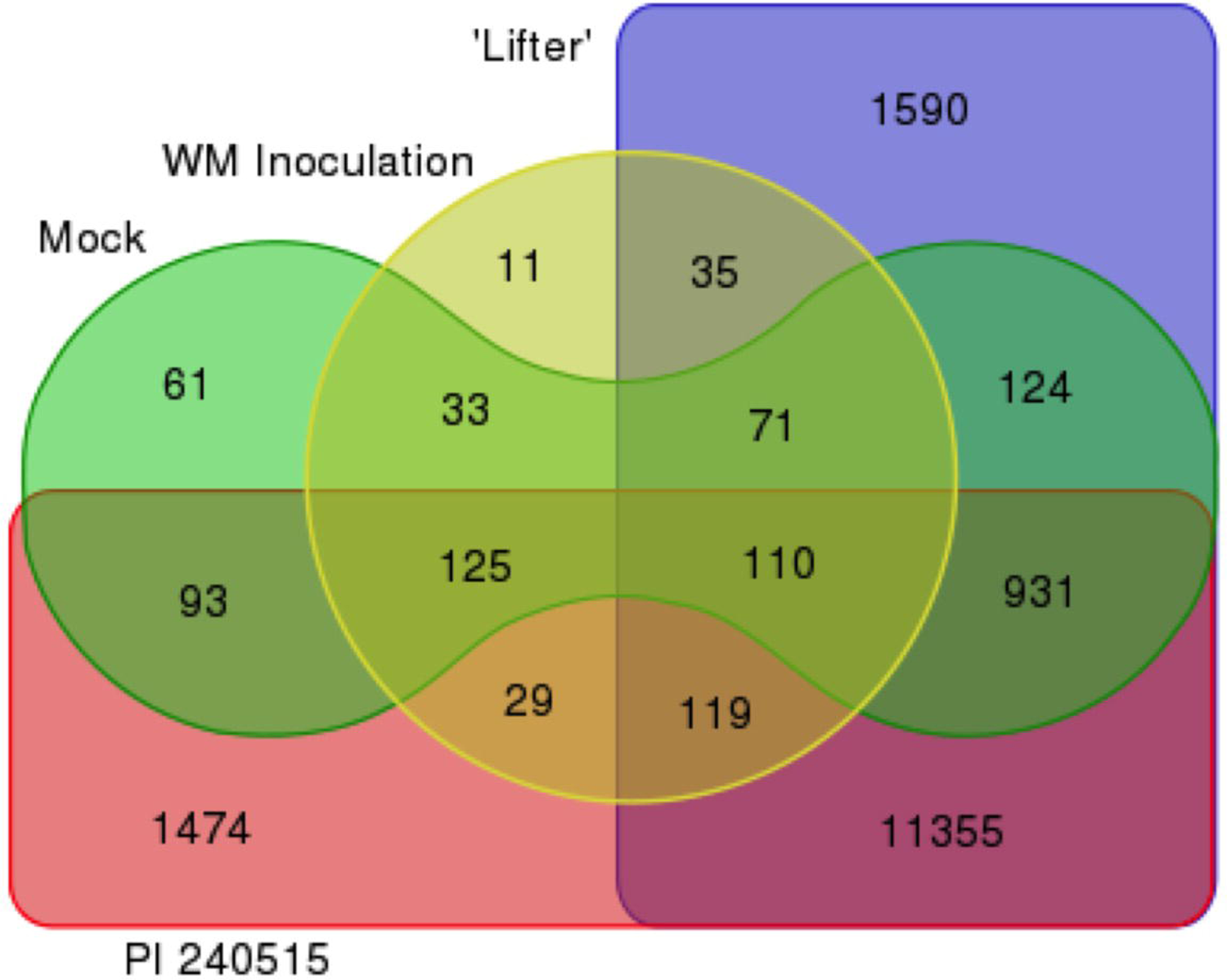
Venn diagram comparisons of time series differential expression analyses. In green, transcripts with significant DE between ‘Lifter’ and PI 240515 in mock samples. In yellow, transcripts with significant DE between ‘Lifter’ and PI 240515 in *S. sclerotiorum* inoculated samples. In purple, transcripts of ‘Lifter’ with significant DE between mock samples and *S. sclerotiorum*-inoculated samples. In pink, transcripts of PI 240515 with significant DE between mock samples and *S. sclerotiorum*- inoculated samples. To narrow the candidate resistance genes pool from all DE, two assumptions were made: (i) a candidate gene should respond to *S. sclerotiorum* inoculation, and (ii) the expression of a candidate gene should have up-regulated and significant DE in the *S. sclerotiorum*-inoculated PI 240515 compared to the *S. sclerotiorum*-inoculated ‘Lifter’ samples, but not the mock samples of PI 240515 compared to the mock samples of ‘Lifter’.

**FIGURE 6.**
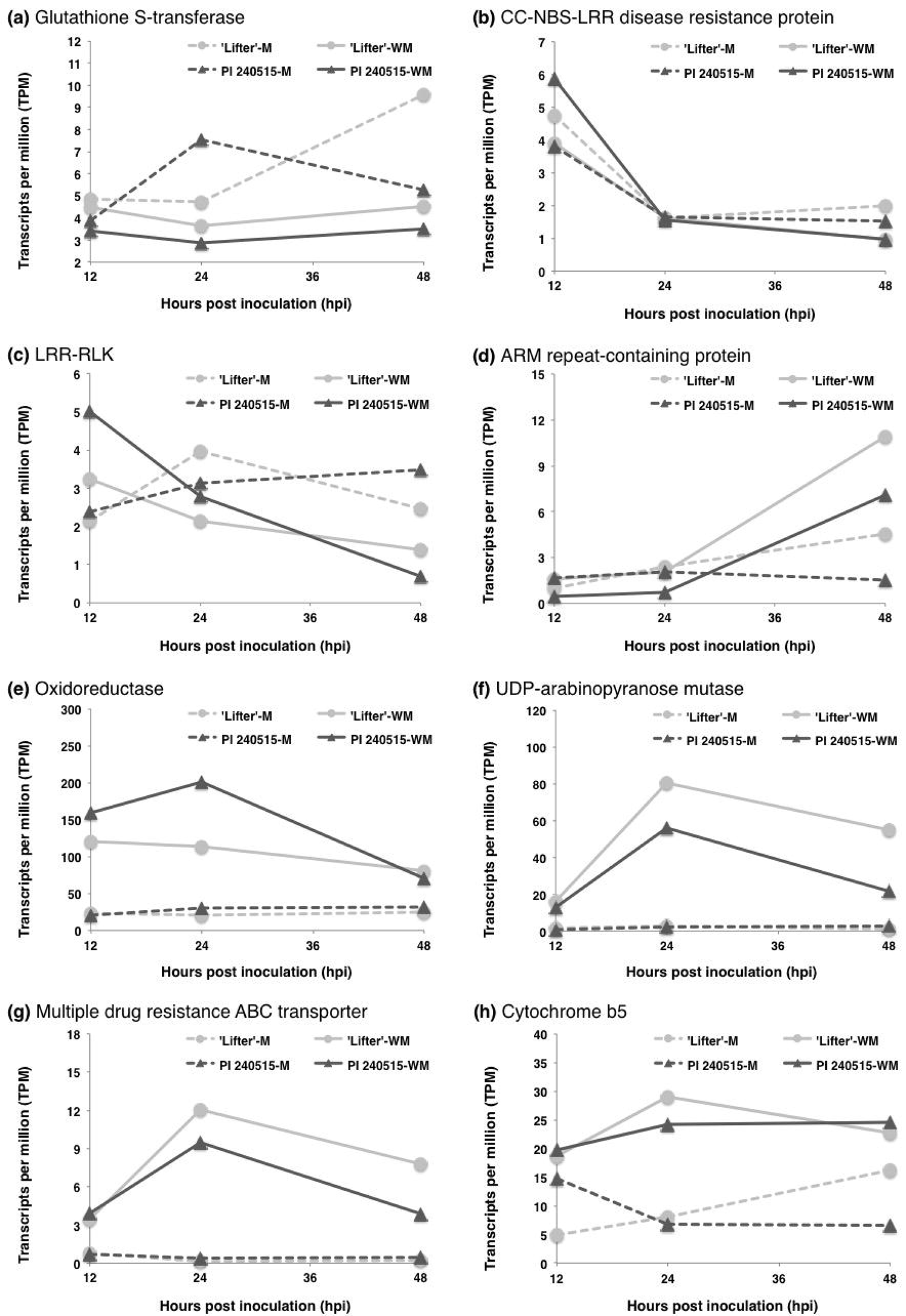
Time course expressions of LRR-RLK transcripts identified from DE analyses. M represents mock samples, and WM represents *S. sclerotiorum*-inoculated samples. (a) TRINITY_DN22904_c0_g1_i2 (b) TRINITY_DN23231_c0_g2_i2 (c) TRINITY_DN18054_c0_g1_i1 (d) TRINITY_DN4777_c0_g1_i1 (e) TRINITY_DN21848_c0_g1_i1.

### 3.7 Integration of GWAS and RNA-Seq results

Integration of results from DE analyses and GWAS identified additional candidate resistance genes, however, most transcripts were down-regulated after *S. sclerotiorum* inoculation (Fig. 3), and only a few transcripts had significantly higher expression in PI 240515 than ‘Lifter’ (Fig. 7, 8). The transcript (TRINITY_DN7903_c0_g1_i2) found for both lesion and nodal resistance, which encodes a putative GST, had differential expression after *S. sclerotiorum* inoculation (Fig. 7a). There were two LRR-containing DE transcripts (TRINITY_DN11274_c0_g2_i1 and TRINITY_DN21987_c1_g2_i1) that significantly associated with lesion resistance (Table 2; Fig. 7b,c). Additionally, five DE transcripts annotated as an ARM repeat superfamily protein (TRINITY_DN21727_c0_g1_i1), an oxidoreductase (TRINITY_DN15345_c0_g1_i2), a UDP-arabinopyranose mutase (TRINITY_DN12885_c0_g1_i1), a multiple drug resistance ABC transporter (TRINITY_DN23674_c1_g2_i1), and a cytochrome b5 (TRINITY_DN29578_c0_g1_i1), were all significantly associated with lesion resistance (Table 2; Fig. 7d-g). On the other hand, there were five DE transcripts annotated as an ACT domain repeat protein (TRINITY_DN5298_c0_g1_i1), a VQ motif-containing protein (TRINITY_DN25769_c0_g1_i1), a β-glucosidase (TRINITY_DN23515_c1_g1_i4), a myoinositol oxygenase (TRINITY_DN21524_c0_g1_i1), and a cytochrome b-561 (TRINITY_DN16214_c1_g2_i1) that were significantly associated with nodal resistance (Fig. 8a-e). Among these transcripts, a putative coiled-coil nucleotide-binding site leucine rich repeat (CC-NBS-LRR) protein appeared to be the most interesting as a lesion resistance candidate because its expression was up-regulated in PI 240515 but down-regulated in ‘Lifter’ after 12 hpi (Fig. 7b). As for nodal resistance, only the putative cytochrome b-561 had higher expression in PI 240515 than ‘Lifter’, and other transcripts mostly had higher DE in ‘Lifter’ than PI 240515 (Fig. 8e).

**FIGURE 7.**
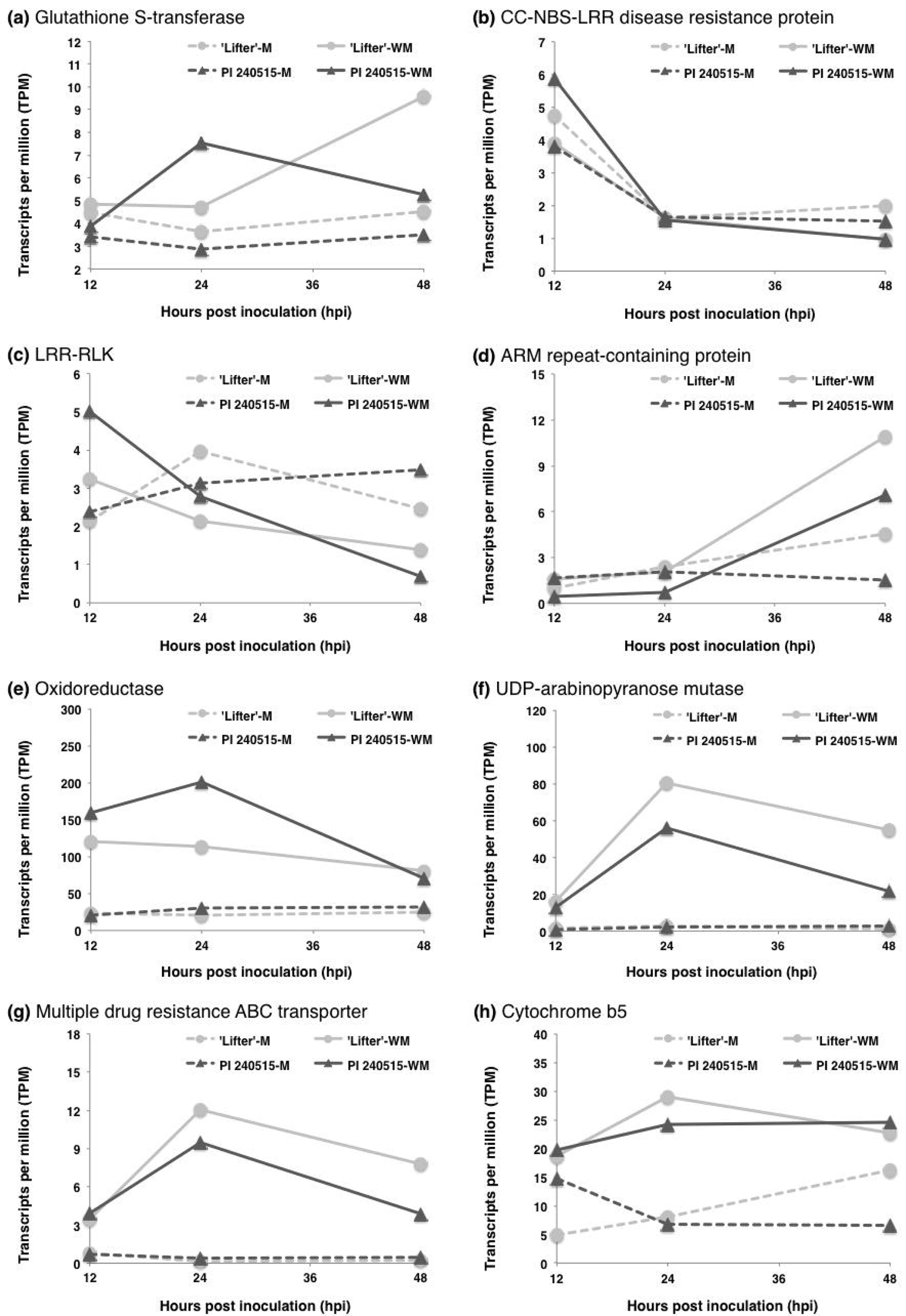
Time course expressions of candidate lesion resistance transcripts identified from both DE analyses and GWAS. M represents mock samples, and WM represents *S. sclerotiorum*-inoculated samples. (a) TRINITY_DN7903_c0_g1_i2 (b) TRINITY_DN11274_c0_g2_i1 (c) TRINITY_DN21987_c1_g2_i1 (d) TRINITY_DN21727_c0_g1_i1 (e) TRINITY_DN15345_c0_g1_i2 (f) TRINITY_DN12885_c0_g1_i1 (g) TRINITY_DN23674_c1_g2_i1 (h) TRINITY_DN29578_c0_g1_i1.

**FIGURE 8.**
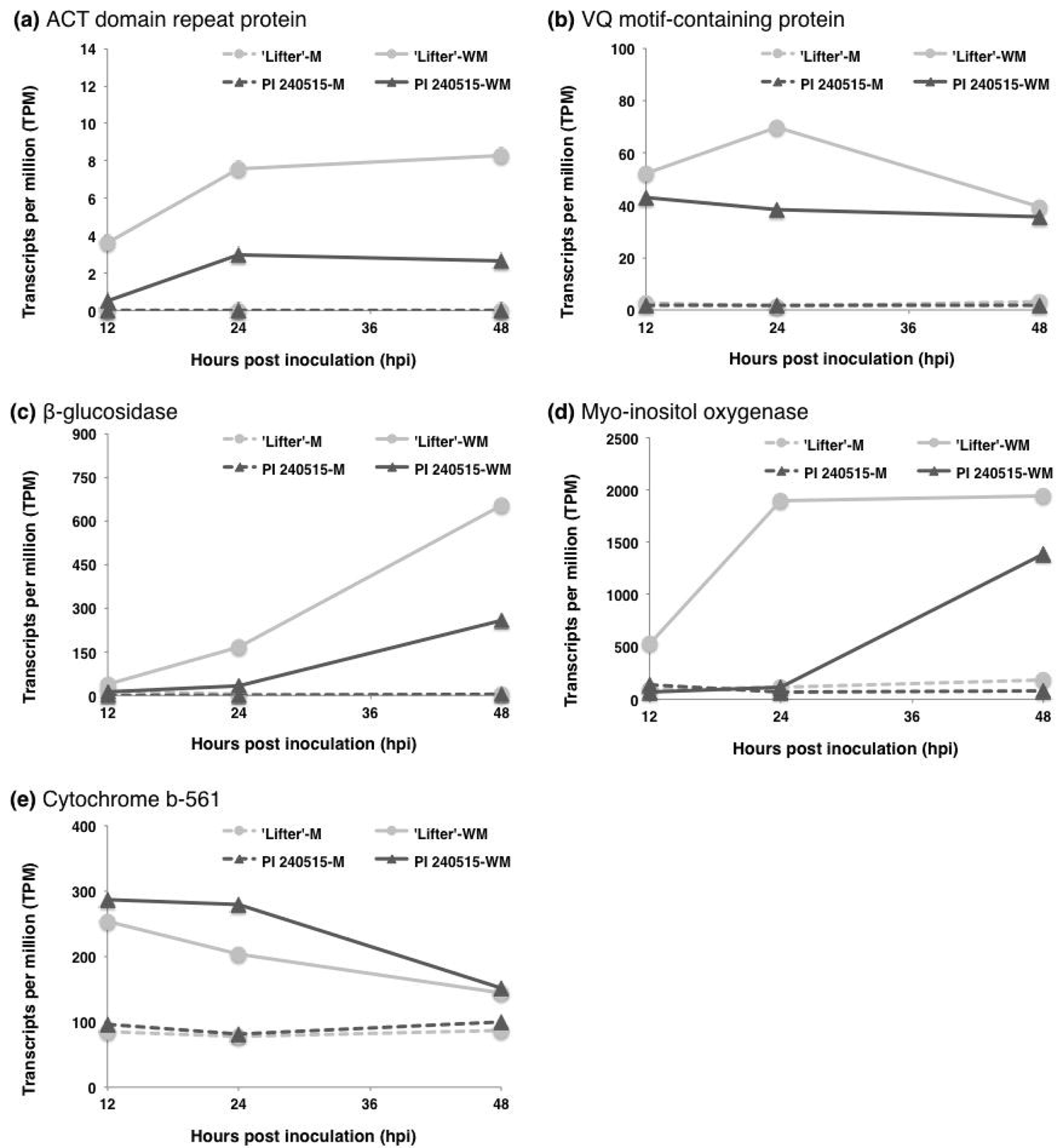
Time course expressions of candidate nodal resistance transcripts identified from both DE analyses and GWAS. M represents mock samples, and WM represents *S. sclerotiorum*-inoculated samples. (a) TRINITY_DN5298_c0_g1_i1 (b) TRINITY_DN25769_c0_g1_i1 (c) TRINITY_DN23515_c1_g1_i4 (d) TRINITY_DN21524_c0_g1_i1 (e) TRINITY_DN16214_c1_g2_i1.

## 4 DISCUSSION

### 4.1 Redox homeostasis is important for both lesion and nodal resistance

In this study, we aimed to understand the genetic makeup of lesion and nodal resistance in pea for resistance to *S. sclerotiorum*. Although there was a weak phenotypic correlation between stem lesion and nodal resistance, GWAS identified different SNPs associated with lesion and nodal resistance ratings. Among hundreds of SNPs, there was only one SNP (C/T), TP13557, found for both phenotypic ratings. The SNP was mapped to a transcript that encodes for a putative glutathione S-transferase (GST) of pea. Interestingly, GST of corn (*Zea mays*) has been identified as a pleiotropic resistance gene for three fungal diseases, southern leaf blight (caused by *Cochliobolus heterostrophus*), gray leaf spot (caused by *Cercospora* species), and northern leaf blight (caused by *Setosphaeria turcica*) in a multivariate mapping study (Wisser et al., 2011). While the significant SNP located in the 3’-UTR of corn GST, three nonsynonymous substitutions were found in the coding sequence, and one of them (histidine to aspartic acid) may contribute about 6% of resistance (Wisser et al., 2011). Other studies also pointed out the importance of GST in potato, rice, and tobacco to *Phytophthora infestans, Magnaporthe oryzae*, and two *Colletotrichum* species, respectively (Dean, Goodwin, & Hisang, 2005; Leonards-Schippers et al., 1994; Wisser et al., 2005). Moreover, most studies focusing on *B. napus* resistance to *S. sclerotiorum* also identified GST regardless of the methodologies, GWAS or RNA-Seq (Girard et al., 2017; Wei et al. 2016; Wu et al., 2016a). A recent GWAS for soybean resistance to *S. sclerotiorum* also identified soybean GST (Wei et al. 2017) which was also noted to have high expression in a transcriptomic study (Calla et al. 2014). Consistently, our results also pointed out GST of pea may play a fundamental role in lesion and nodal resistance to *S. sclerotiorum*. GST has diverse molecular functions in a cell to balance redox homeostasis, and glutathione is important for maintaining a reducing status for cell survival (Tew, 2007). Accordingly, GST may be involved in prohibiting the switch from hemibiotrophic to necrotrophic stage of *S. sclerotiorum*. Another GST function is to detoxify phytotoxins and oxidative substances such as ROS (Wisser et al., 2011), and these functions may slow *S. sclerotiorum* infection and plant cell death. Other than GST, redox-related genes were up-regulated after *S. sclerotiorum* inoculation and were significantly enriched in cluster IV, and redox-related enzymes such as an oxidoreductase and a cytochrome b-561 were found in GWAS for lesion or nodal resistance, respectively. Although the molecular function of cytochrome b-561 in plant resistance is not clear, it has been also discovered for *S. sclerotiorum* resistance in *B. napus* using GWAS and RNA-Seq (Wei et al., 2016). Surprisingly, our combined RNA-Seq and GWAS strategy to search for *S. sclerotiorum* resistance in pea ended up with results similar to the study of *B. napus*, where β-glucosidase, TPR-containing protein, ARM repeat superfamily protein, cytochrome b-561, LRR-containing proteins, and the GST were also found for *B. napus* (Table S2, S3; Wei et al., 2016). Together, our results support the importance of homeostasis for *S. sclerotiorum* resistance, and we identified many potential redox-related transcripts as well as others with roles in basal resistance to white mold.

### 4.2 Lesion resistance

Several LRR-containing genes were found in transcriptomic studies of *B. napus*-*S. sclerotiorum* (Wei et al., 2016; Wu et al., 2016b). The results of our study also discovered several up-regulated LRR-containing transcripts for lesion resistance but not nodal resistance. Two transcripts with evidence from both GWAS and RNA-Seq are a putative CC-NBS-LRR protein and an LRR-RLK protein (Fig. 7b,c). Both transcripts had higher expression in PI 240515 at 12 hpi but the expressions dropped over time to the expression level of ‘Lifter’. Although it is well known that LRR-containing proteins contribute to R-gene based resistance in plants to biotrophic pathogens (Kushalappa, Yogendra, & Karre, 2016), it is unclear how much these LRR-containing transcripts are involved in lesion resistance to *S. sclerotiorum*. Moreover, these LRR-containing transcripts were not discovered in GWAS for nodal resistance, which indicated the possibility that the lesion resistance relies on LRR-containing proteins more than nodal resistance. Other than the LRR type of tandem repeats, several ARM repeat-containing proteins were found for lesion resistance by GWAS (Table 2, Table S2). It has been shown that an ARM-containing ligase in rice negatively controls resistance (Li et al., 2012; Sharma & Pandey, 2015), indicating higher expression in ‘Lifter’ might favor *S. sclerotiorum* infection (Fig. 7d). While plant cell wall synthesis enzymes such as cellulose synthase were identified, only a putative UDP-arabinopyranose mutase was up-regulated after *S. sclerotiorum* inoculation (Fig. 7f). Similarly, two pleiotropic drug resistance ABC transporters were found (Table 2) but only one displayed up-regulation after *S. sclerotiorum* inoculation (Fig. 7g). It has been shown that a pleiotropic drug resistance ABC transporter is involved in resistance to *Botrytis cinerea*, a closely-related fungal species to *S. sclerotiorum* (Stukkens et al., 2005). Accordingly, our results suggested diverse mechanisms were involved in lesion resistance to limit *S. sclerotiorum* expansion.

### 4.3 Nodal resistance

Five transcripts found by both GWAS and RNA-Seq had higher DE after *S. sclerotiorum* inoculation (Fig. 7). While the ACT domain repeat-containing proteins have diverse functions in plant physiologies (Feller, Yuan, & Grotewold, 2017), the β-glucosidase might be involved in cell wall reinforcement or releasing damage associated molecular patterns (DAMP) (Duran-Flires & Heil, 2016). Additionally, one of the transcripts identified for nodal resistance is a putative VQ motif-containing protein, which had higher expression in ‘Lifter’ than PI 240515 at 24 hpi (Fig. 8b). It has been shown that two VQ motif-containing proteins, VQ12 and VQ29, in *Arabidopsis* negatively regulate resistance to *B. cinerea*. Down regulation of VQ12 and VQ29 by miRNA silencing promoted *Arabidopsis* resistance to *B. cinerea* while overexpression increased susceptibility (Wang et al., 2015). In addition, overexpression of *Arabidopsis* VQ5 and VQ20 demonstrated enhanced susceptibility to *B. cinerea* and *Pseudomonas syringae* (Cheng et al., 2012). These discoveries might help to explain the potential functions of this VQ motif-containing protein in pea and higher expression in ‘Lifter’ at 24 hpi potentially favoring *S. sclerotiorum* infection.

Another transcript with earlier and higher expression in ‘Lifter’ was a putative myo-inositol oxygenase (Fig. 7d). Myo-inositol is a product catalyzed from glucose-6-phosphate by the myo-inositol-1-phosphate synthase, and it can be further metabolized into UDP-glucuronic acid by myo-inositol oxygenase (Kanter et al., 2005). One of the functions of UDP-glucuronic acid is being the precursor of plant cell wall polysaccharides, and under the circumstance, earlier and higher expression of this transcript in ‘Lifter’ may indicate the need of plant cell wall reinforcement under high *S. sclerotiorum* pressure. Gene expression difference for myo-inositol metabolism was also reported in resistant and susceptible soybeans to *S. sclerotiorum* (Calla et al., 2009). Additionally, myo-inositol is also the precursor of galactinol, which has been suggested to induce resistance against syncytia development for the cyst nematode *Heterodera schachtii* (Siddeigue et al., 2014). When myo-inositol oxygenase processes myo-inositol into UDP-glucuronic acid, the metabolism bypasses and reduces the production of galactinol for inducing disease resistance signaling (Cho et al., 2010; Kim et al., 2008). Under this scenario, disease progress in ‘Lifter’ could be faster than in PI 240515. Because the expression patterns of nodal resistance-related transcripts generally had a delay in PI 240515 compared to ‘Lifter’ and because *S. sclerotiorum* can still infect PI 240515 albeit at a slower rate than ‘Lifter’, it is possible that down-regulation of transcripts favoring *S. sclerotiorum* infection underlies nodal resistance in PI 240515. More studies are needed to reveal the molecular mechanisms of nodal resistance.

### 4.4. Summary

In this study, we applied GWAS and RNA-Seq to understand lesion and nodal resistance in pea to *S. sclerotiorum*. Other than a transcript encoding GST, which was discovered for both lesion and nodal resistance, our results pointed out different mechanisms underlying lesion and nodal resistance. We revealed SNPs exclusively for the lesion and nodal resistance, and together with time series DE analyses, we suggested these two types of white mold resistance are differently controlled by a diverse of genetic mechanisms.

## ACKNOWLEDGEMENTS

This study was supported by the National Sclerotinia Initiative, grant ID: 58-5442-9-239, USDA-ARS. We thank Patricia Santos and J. Alejandro Rojas for helpful discussions in preparation of this manuscript.

## ACCESSION NUMBER

The Illumina sequences were deposited at SRA database under BioProject accession number PRJNA261444.

## AUTHOR CONTRIBUTION

H.X.C., H.S., X.Z., and L.P. conducted plant inoculation and phenotyping. H.X.C, H.S., and J.W. performed data analyses. K.M., L.P., and M.I.C. maintained plant and fungal materials. X.Z. generated the RNA-Seq data. H.X.C. and M.I.C. led the manuscript writing, and all the co-authors refined and approved the manuscript.

## SUPPLEMENTARY TABLES

**TABLE S1** Illumina sequencing statistics

**TABLE S2** Significant SNPs associated with lesion resistance

**TABLE S3** Significant SNPs associated with nodal resistance

**TABLE S4** Significant DEs of the 119 candidate transcripts controlling *S. sclerotiorum* resistance and susceptibility

**TABLE S5** Significant DEs of the 135 candidate transcripts controlling *S. sclerotiorum* resistance and susceptibility

**FIGURE S1** Gene Ontology (GO) using singular enrichment analysis for transcripts in the cluster I. (a) Biological process. (b) Cellular component. (c) Molecular function.

**FIGURE S2** Gene Ontology (GO) using singular enrichment analysis for transcripts in the cluster II. (a) Biological process. (b) Cellular component. (c) Molecular function.

**FIGURE S3** Gene Ontology (GO) using singular enrichment analysis for transcripts in the cluster III. Only some GO terms in the cellular component category were significant.

## REFERENCES

Andrews, S. (2010). FastQC: a quality control tool for high throughput sequence data. Available online at:http://www.bioinformatics.babraham.ac.uk/projects/fastqc

Bastien, M., Sonah, H., & Belzile, F. (2014). Genome wide association mapping of *Sclerotinia sclerotiorum* resistance in soybean with a genotyping-by-sequencing approach. Plant Genome, 7, 1-13. doi:10.3835/plantgenome2013.10.0030

Bolger, A.M., Lohse, M., & Usadel, B. (2014). Trimmomatic: a flexible trimmer for Illumina sequence data. Bioinformatics, 30, 2114-2120. doi:10.1093/bioinformatics/btu170

Bolton, M.D., Thomma, B.P.H.J., & Nelson, B.D. (2006). *Sclerotinia sclerotiorum* (Lib.) de Bary: biology and molecular traits of a cosmopolitan pathogen. Molecular Plant Pathology 7, 1-16. doi: 10.1111/J.1364-3703.2005.00316.X

Bray, N.L., Pimentel, H., Melsted, P., & Pachter, L. (2017). Near-optimal probabilistic RNA-Seq quantification. Nature Biotechnology, 34, 525-527.

Bush, W.S., & Moore, J.H. (2012). Chapter 11: genome-wide association studies. PloS Computational Biology, 8, e1002822. doi:10.1371/journal.pcbi.1002822

Calla, B., Voung, T., Radwan, O., Hartman, G.L., & Clough, S.J. (2009). Gene expression profiling soybean stem tissue early response to *Sclerotinia sclerotiorum* and *in silico* mapping in relation to resistance markers. Plant Genome, 2, 149-166. doi:10.3835/plantgenome2008.02.0008

Calla, B., Blahut-beatty, L., Koziol, L., Simmonds, D.H., & Clough, S.J. (2014). Transcriptome analyses suggest a disturbance of iron homeostasis in soybean leaves during white mold disease establishment. Molecular Plant Pathology, 15, 576-588. doi: 10.1111/mpp.12113

Cheng, Y., Zhou, Y., Yang, Y., Chi, Y.-J., Zhou, J., Chen, J.-Y., … Chen, Z. (2012). Structural and functional analysis of VQ motif-containing proteins in Arabidopsis as interacting proteins of WRKY transcription factors. Plant Physiology, 159, 810-825. doi: 10.1104/pp.112.196816

Cho, S.M., Kang, E.Y., Kim, M.S., Yoo, S.J., Im, Y.J., Kim, Y.C., … Cho, B.H. (2010). Jasmonate-dependent expression of a galactinol synthase gene is involved in priming of systemic fungal resistance in *Arabidopsis thaliana*. Botany, 88, 452-461. doi: 10.1139/B10-009

Collier, S.M., & Moffett, P. (2009). NB-LRRs work a “bait and switch” on pathogens. Trends in Plant Science, 14, 521-529. doi: 10.1016/j.tplants.2009.08.001

Dean, J.D., Goodwin, P.H., & Hisang, T. (2005). Induction of glutathion S-transferase genes of Nicotiana benthamiana following infection by *Colletotrichum destructivum* and *C. orbiculare* and involvement of one in resistance. Journal of Experimental Botany, 56, 1525-1533. doi: 10.1093/jxb/eri145

Derbyshire, M., Denton-Giles, M., Hegedus, D., Seifbarghi, S., Rolins, J., van Kan, J., … Oliver, R. (2017). The complete genome sequence of phytopathogenic fungus *Sclerotinia sclerotiorum* reveals insights into the genome architecture of broad host range pathogens. Genome Biology and Evolution, 9, 593-618. doi:10.1093/gbe/evx030

Duran-Flires, D., & Heil, M. (2016). Sources of specificity in plant damaged-self recognition. Current Opinion in Plant Biology, 32, 77-87. doi: 10.1016/j.pbi.2016.06.019

Feller, A, Yuan, L., & Grotewold, E. (2017). The BID domain in plant bHLH proteins is like an ACT-like domain. Plant Cell, 29, 1800-1802. doi: 10.1105/tpc.17.00356

Gordon, A. (2014). FASTX-toolkit. Available online at http://hannonlab.cshl.edu/fastx_toolkit/index.html

Grabherr, M.G., Haas, B.J., Yassour, M., Levin, J.Z., Thompson, D.A., Amit, I., … Regev A. (2011). Full-length transcriptome assembly from RNA-seq data without a reference genome. Nature Biotechnology, 29, 644-52. doi: 10.1038/nbt.1883

Girard, I.J., Tong, C., Becker, M.G., Mao, X., Huang, J., de Kievit, T. … Belmonte, M.F. (2017). RNA sequencing of Brassica napus reveals cellular redox control of *Sclerotinia* infection. Journal of Experimental Botany, erx338. doi: 10.1093/jxb/erx338

Haas, B.J., Papanicolaou, A., Yassour, M., Grabherr, M., Blood, P.D., Bowden, J., … Regev A. (2013). De novo transcript sequence reconstruction from RNA-seq using the Trinity platform for reference generation and analysis. Nature Protocol, 8, 1494-512. doi: 10.1038/nprot.2013.084

Hobbs, T.Q., Schmitthenner, A.E., & Ellett, C.W. (1981) Top dieback of soybean caused by *Diaporthe phaseolorum* var. *caulivora*. Plant Disease, 65, 618-620.

Holdsworth, W.L., Gazacve, E., Cheng, P., Myers, J.R., Gore, M.A., Coyne, C.J., … Mazourek, M. (2017). A community resource for exploring and utilizing genetic diversity in the USDA pea single plant plus collection. Horticulture Research, 4, 17017. doi:10.1038/hortres.2017.17

Kabbage, M., Yarden, O., & Dickman, M.B. (2015). Pathogenic attributes of *Sclerotinia sclerotiorum*: switching from a biotrophic to necrotrophic lifestyle. Plant Science, 233, 53-60. doi: 10.1016/j.plantsci.2014.12.018

Kanter, U., Usadel, B., Guerineau, F., Li, Y., Pauly, M., & Tenhaken, R. (2005). The inositol oxygenase gene family of *Arabidopsis* is involved in the biosynthesis of nucleotide sugar precursors for cell-wall matrix polysaccharides. Planta, 221, 243-254. doi: 10.1007/s00425-004-1441-0

Kerr, S.C., Gaiti, F., Beveridge, C.A., & Tanurdzic, M. (2017). De novo transcriptome assembly reveals high transcriptional complexity in *Pisum sativum* axillary buds and shows rapid changes in expression of diurnally regulated genes. BMC Genomics, 18, 221. doi: 10.1186/s12864-017-3577-x

Kim, D., Pertea, G., Trapnell, C., Pimental, H., Kelly, R., & Salzberg, S.L. (2013). TopHat2: accurate alignment of transcriptomes in the presence of insertions, deletions and gene fusions. Genome Biology, 14, R36. doi: 10.1186/gb-2013-14-4-r36

Kim, M.S., Cho, S.M., Kang, E.Y., Im, Y.J., Hwangbo, H., Kim, Y.C., … Cho, B.H. (2008). Galactinol is a signaling component of the induced systemic resistance caused by *Pseudomonas chlororaphis* O6 root colonization. Molecular Plant-Microbe Interactions, 21, 1643-1653. doi: 10.1094/MPMI-21-12-1643

Kushalappa, A.C., Yogendra, K.N., & Karre, S. (2016). Plant innate immune response: qualitative and quantitative resistance. Critical Reviews in Plant Sciences, 35, 38-55. doi: 10.1080/07352689.2016.1148980

Leonards-Schippers, C., Gieffers, W., Schafer-Pregl, R., Ritte, E., Knapp, S.J., Salamini, F., & Gebhardt, C. (1994). Quantitative resistance to *Phytophthora infestans* in potato: a case study for QTL mapping in an allogamous plant species. Genetics, 137, 67-77.

Li, W., Ahn, I., Ning, Y., Park, C.-H., Zeng, L., Whitehill, J.G.A., … Wang, G.-L. (2012). The U-box/ARM E3 ligase PUB13 regulates cell death, defense, and flowering time in Arabidopsis. Plant Physiology, 159, 239-250. doi: 10.1104/pp.111.192617

Lu, K., Peng, L., Zhang, C. Lu, J., Yang, B., Xiao, Z., …, Li, Jiana. (2017). Genome-wie association and transcriptome analyses reveal candidate genes underlying yield-determining traits in Brassica napus. Frontiers in Plant Science, 8, 206. doi: 10.3389/fpls.2017.00206

Mbengue, M., Navaud, O., Peyraud, R., Barascud, M., Badet, T., Vincent, R., …, Raffaela, S. (2016) Emerging trends in molecular interactions between plants and the broad host range fungal pathogens *Botrytis cinerea* and *Sclerotinia sclerotiorum*. Frontiers in Plant Science, 7, 422. doi: 10.3389/fpls.2016.00422

McCaghey, M., Willbur, J., Ranjan, A., Grau, C.R., Chapman, S., Diers, B., …, Smith, D.L. (2017). Development and evaluation of *Glycine max* germplasm lines with quantitative resistance to *Sclerotinia sclerotiorum*. Frontiers in Plant Science, 8, 1495. doi: 10.3389/fpls.2017.01495

McPhee K. E., & Muehlbauer, F. J. (2002). Registration of ‘LIFTER’ green dry pea. Crop Science, 42, 1377–1378.

Moellers, T.C., Singh, A., Zhang, J., Brungardt, J., Kabbage, M., Meuller, D.S., … Singh, A.K. (2017). Main and epistatic loci studies in soybean for *Sclerotinia sclerotiorum* resistance reveal multiple modes of resistance in multi-environments. Scientific Reports, 7, 3554. doi:10.1038/s41598-017-03695-9

Muchero, W., Ehlers, J.D., Close, T.J., & Roberts, P.A. (2011) Genic SNP markers and legume synteny reveal candidate genes underlying QTL for *Macrophomina phaseolina* resistance and maturity in cowpea [*Vigna unguiculata* (L) Walp.]. BMC Genomics, 12, 8.

Murtagh, F., & Legendre, P. (2014). Ward’s hierarchical agglomerative clustering method: which algorithms implement Ward’s criterion? Journal of Classification, 31, 274-295. doi: 10.1007/s00357-014-9161-z

Peltier, A. J., Hatfield, R. D., & Grau, C. R. (2009). Soybean stem lignin concentration relates to resistance to *Sclerotinia sclerotiorum*. Plant Disease, 93, 149-154. doi: 10.1094/PDIS-93-2-0149

Porter, L.D., Hoheisel, G., & Coffman, V.A. (2009). Resistance of peas to *Sclerotinia sclerotiorum* in the *Pisum* core collection. Plant Pathology, 58, 52-60. doi: 10.1111/j.1365-3059.2008.01937.x

Porter, L. (2011). Selection of pea genotypes with partial resistance to *Sclerotinia sclerotiorum* across a wide range of temperatures and periods of high relative humidity. Euphytica, 186, 671-678. doi: 10.1007/s10681-011-0531-x

Purcell, S., Neale, B., Todd-Brown, K., Thomas, L., Ferreira, M.A.R., Bender, D., …, Sham, P.C. (2007). PLINK: a tool set for whole-genome association and population-based linkage analyses. The American Journal of Human Genetics, 81, 559-575. doi: 10.1086/519795

Seifbarghi, S., Borhan, M.H., Wei, Y., Coutu, C., Robinson, S.J., & Hegedus, D.D. (2017). Changes in the *Sclerotinia sclerotiorum* transcriptome during infection of Brassica napus. BMC Genomics, 18, 266. doi: 10.1186/s12864-017-3642-5

Sharma, M., & Pandey, G.K. (2015). Expansion and function of repeat domain proteins during stress and development in plants. Frontier in Plant Science, 6, 1218. doi: 10.3389/fpls.2015.01218

Siddeigue, S., Endres, S., Sobczak, M., Radakovic Z.S., Fragner, L., Florian, M.W., … Bohlmann, H. (2014). Myo-inositol oxygenase is important for the removal of excess myoinositol from syncytia induced by *Heterodera schachtii* in Arabidopsis roots. New Phytologist, 201, 476-485. doi: 10.1111/nph.12535

Stukkens, Y., Bultreays, A., Grec, S., Trombik, T., Vangam, D., & Boutry, M. (2005). NpPDR1, a pleiotropic drug resistance-type ATP-binding cassette transporter from *Nicotiana plumbaginifolia*, plays a major role in plant pathogen defense. Plant Physiology, 139, 341-351. doi: 10.1104/pp.105.062372

Tayeh, N., Aubert, G., Oilet-Nayel, M., Lejeune-Henaut, I., Warkentin, T.D., & Burstin, J. (2015). Genomic tools in pea breeding programs: status and perspectives. Frontiers in Plant Science, 6, 1037. doi: 10.3389/fpls.2015.01037

Tew, K.D. (2007). Redox in redux: emergent roles for glutathione S-transferase P (GSTP) in regulation of cell signaling and S-glutathionylation. Biochemical pharmacology, 73, 1257-1269. doi:10.1016/j.bcp.2006.09.027

Tian, T., Liu, Y., Yan, H., You, Q., Yi, X., Du, Z., … Su, Z. (2017) agriGO v2.0: a GO analysis toolkit for the agricultural community, 2017 update. Nucleic Acid Research, 45, W122-W129. doi: 10.1093/nar/gkx382.

Wang, H., Hu, Y., Pan, J., & Yu, D. (2015). Arabidopsis VQ motif-containing proteins VQ12 and VQ29 negatively modulate basal defense against Botrytis cinerea. Scientific Reports, 5, 14185. doi: 10.1038/srep14185

Wei, L., Jian, H., Lu, K., Filardo, F., Yin, N., Liu, L., … Li, J. (2016). Genome-wide association analysis and differential expression analysis of resistance to *Sclerotinia* stem rot in *Brassica napus*. Plant Biotechnology Journal, 14, 1368-1380. doi: 10.1111/pbi.12501

Wei, W., Mesquita, A.C.O., Figueiró, A.D.A., Wu, X., Manjunatha, S., Wickland, D.P.,…Clough, S.J. (2017) Genome-wide association mapping of resistance to a Brazilian isolate of *Sclerotinia sclerotiorum* in soybean genotypes mostly from Brazil. BMC Genomics, 18, 849. doi: 10.1186/s12864-017-4160-1

Wei, W., & Clough, S.J. (2016) *Sclerotinia sclerotiorum*: molecular aspects in plant-pathogenic interactions. Revisao Anual De Patologia De Plantas, 24, 174-189.

Williams, B., Kabbage, M., Kim, H.-J., Britt, R., and Dickman, M.B. (2011). Tipping the balance: *Sclerotinia sclerotiorum* secreted oxalic acid suppresses host defenses by manipulating the host redox environment. PLoS Pathogens, 7, e1002107. doi: 10.1371/journal.ppat.1002107

Wisser, R.J., Kolkman, J.M., Patzoldt, M.E., Holland, J.B., Yu, J., Krakowsky, M., … Balunt-Kurti, P.J. (2011). Multivariate analysis of maize disease resistances suggests a pleiotrophic genetic basis and implicates a GST gene. Proceedings of the National Academy of Sciences, 108, 7338-7344. doi: 10.1073/pnas.1011739108

Wisser, R.J., Sun, Q., Hukbert, S.H., Kresovich, S., & Nelson, R.J. (2005). Identification and characterization of regions of the rice genome associated with broad-spectrum, quantitative disease resistance. Genetics, 169, 2277-2293. doi: 10.1534/genetics.104.036327

Wu, J., Zhao, Q., Liu, S., Shahid, M., Lan, L., Cai, G., …, Zhou, Y. (2016a). Genomie-wide association study identifies new loci for resistance to *Sclerotinia* stem rot in *Brassica napus*. Frontiers in Plant Science, 7, 1418. doi: 10.3389/fpls.2016.01418

Wu, J., Zhao, Q., Yang, Q., Liu, H., Li, Q., Yi, X., …, Zhou, Y. (2016b). Comparative transcriptomic analysis uncovers the complex genetic network for resistance to *Sclerotinia sclerotiorum* in *Brassica napus*. Scientific Reports, 6, 19007. doi: 10.1038/srep19007

Xu, L., Xiang, M., White, D., & Chen, W. (2015). pH dependency of sclerotial development and pathogenicity revealed by using genetically defined oxalate-minus mutants of *Sclerotinia sclerotiorum*. Environmental Microbiology, 17, 2896-2909. doi: 10.1111/1462-2920.12818

Zhou, J., Sun, A., & Xing, D. (2013). NADPH oxidase confers *Arabidopsis* resistance to *Sclerotinia sclerotiorum*. Journal of Experimental Botany, 64, 3261-3272. doi:10.1093/jxb/ert166

Zhuang, X., McPhee, K.E., Coram, T.E., Peever, T.L., & Chilvers, M.I. (2012). Rapid transcriptome characterization and parsing of sequences in a non-model host-pathogen interaction: pea-*Sclerotinia sclerotiorum*. BMC Genomics, 13, 668. doi: 10.1186/1471-2164-13-668

Zhuang, X., McPhee, K.E., Coram, T.E., Peever, T.L., & Chilvers, M.I. (2013). Development and characterization of 37 novel EST-SSR markers in *Pisum sativum* (Fabaceae). Applications in Plant Sciences, 1, 1200249. doi:10.3732/apps.1200249

